# Deciphering the transcriptomic regulation of heat stress responses in *Nothofagus pumilio*

**DOI:** 10.1101/2021.01.25.428064

**Authors:** Maximiliano Estravis-Barcala, Katrin Heer, Paula Marchelli, Birgit Ziegenhagen, María Verónica Arana, Nicolás Bellora

## Abstract

Global warming is predicted to exert negative impacts on plant growth due to the damaging effect of high temperatures on plant physiology. Revealing the genetic architecture underlying the heat stress response is therefore crucial for the development of conservation strategies, and for breeding heat-resistant plant genotypes. Here we investigated the transcriptional changes induced by heat in *Nothofagus pumilio*, an emblematic tree species of the sub-Antarctic forests of South America. Through the performance of RNA-seq of leaves of plants exposed to 20°C (control) or 34°C (heat shock), we generated the first transcriptomic resource for the species. We also studied the changes in protein-coding transcripts expression in response to heat. We found 5,214 contigs differentially expressed between temperatures. The heat treatment resulted in a down-regulation of genes related to photosynthesis and carbon metabolism, whereas secondary metabolism, protein re-folding and response to stress were up-regulated. Moreover, several transcription factor families like WRKY or ERF were promoted by heat, alongside spliceosome machinery and hormone signaling pathways. Through a comparative analysis of gene regulation in response to heat in *Arabidopsis thaliana, Populus tomentosa* and *N. pumilio* we provide evidence of the existence of shared molecular features of heat stress responses across angiosperms, and identify genes of potential biotechnological application.

## Introduction

Heat stress is becoming a threat for food security as global warming progresses [1], and has also the potential to affect biodiversity, primary productivity and ecological functions in natural ecosystems [2]. The ability of plants to respond to different climatic scenarios is critical to their long-term persistence in natural habitats. It is thus imperative to comprehend the genetic architecture of the responses of plants to high temperature in order to better anticipate species’ performance under global warming, and for the detection of genotypes with higher ability to grow under these conditions.

Much of our knowledge about the molecular bases of plants’ responses to heat stress is rooted on studies in short-lived plants, such as the model species *Arabidopsis thaliana*, whereas information in trees is generally scarce [3]. In *A. thaliana*, responses to heat stress are governed by complex transcriptional pathways. The Heat Shock Transcription Factor A and B families (HSFA and HSFB respectively) are considered key regulatory components, inducing the transcription of many stress-related genes such as Heat Shock Proteins (HSPs) and ROS scavenger enzymes [4]. Particularly, HSFA1 promotes the activation of transcriptional networks through the regulation of other relevant stress transcription factors such as Dehydration-Responsive Element Binding Protein 2A (DREB2A), which activates heat-responsive genes like HSPs [5]. Studies in the tree model species *Populus trichocarpa* (black cottonwood) indicate that HSF proteins also regulate HSPs expression in trees [6, 7]. In addition to the classical HSFA and B pathways, abscisic acid (ABA) accumulation and signaling is stimulated by heat in *A. thaliana* [8]. ABA activates DREB2A and Abscisic Acid-Responsive Element Binding Protein 1 (AREB1), which act synergistically with HSFA6b in the promotion of the expression of heat stress related genes [9]. However, we still do not know whether the components and signaling pathways described in *A. thaliana* are conserved among plant species, and studies comparing genetic architecture of heat stress among annual and perennial plants are scarce [3].

In the last ten years, Next Generation Sequencing (NGS) techniques revolutionized genomics and allowed in-depth genomic studies of non-model species [10–12].

Particularly, messenger RNA sequencing (RNA-seq), in combination with *de novo* transcriptome assembly, offers a unique opportunity to study gene expression on a global scale related with a given developmental or environmental condition, even in species lacking reference genomes. Notwithstanding this, transcriptomic studies in relation to the responses of heat stress in trees are limited, and mostly involving species of the northern hemisphere [13–18]. The study of heat-mediated gene expression on a global scale in a wide spectrum of forestry species constitutes thus a priority for the understanding of the diversification of molecular strategies that trees evolved to cope with changes in environmental temperature, and to gain insight into their adaptation to the local environment.

The southern region of the Andes hosts rainforest and sub-Antarctic temperate *Nothofagus* forests across a narrow landmass that spans ca. 20° of latitude. These forests embrace an extraordinary ecological diversity across different environments that will be affected by increasing temperatures according to predictions of global climate change [19]. *Nothofagus pumilio* is one of the most widely distributed species of this region and occurs from the northern Patagonian Andes and central Chilean region (35°S) to the high latitudes at Tierra del Fuego (55°S). Thus, it inhabits an iconic latitudinal gradient that denotes strong adaptation to diverse environmental conditions [20, 21]. However, *N. pumilio* shows an unusual dependence of its altitudinal distribution with latitude not found in other native species of the region. It ranges in elevation from 0 to 2000 meters above sea level (m a.s.l), but north of 41°S it grows only in the sub-Alpine colder zone where it commonly forms the treeline. On the other hand, in the southern part of its range, in colder environments of high latitudes, it occurs both at high (treeline) and low (sea level) elevations [21]. This suggests a strong susceptibility of the species to grow in relatively warm environments. Understanding the response of *N. pumilio* to heat stress thus becomes a priority for the development of conservation strategies and the identification of heat-resistant genotypes able to cope with increasing temperatures predicted by global climate change.

In this study, we aimed to gain insight into the genetic architecture of the responses of *N. pumilio* to heat stress and to identify genes that might work as candidates of this response in *N. pumilio*. For this purpose, here we present the first assembled and annotated transcriptome for *N. pumilio*, and investigate differential gene expression in protein-coding transcripts in response to heat. We also compare our results with previously published studies in other plant species in order to help elucidate shared molecular features of plant heat stress response.

## Materials and methods

### Description of the species

*Nothofagus pumilio* belongs to Nothofagaceae (Kuprianova), a monotypical family of deciduous and evergreen trees from the southern hemisphere in the order Fagales, which includes oaks, beeches, chestnuts, alders, birches, hazelnuts, and other well-known trees. It constitutes an iconic species of the South America temperate forests and its distribution spans the narrow forest landmass of the Andes, covering ca. 2500 km in southern-northern direction [20, 21]. Due to the high quality of its wood and its wide distribution, *N. pumilio* constitutes one of the most economically important native species of Patagonia [22].

### Plant material, growth conditions and heat stress treatments

In order to perform heat stress experiments, *N. pumilio* seedlings were grown from seeds collected in Challhuaco, San Carlos de Bariloche, Argentina (latitude: −41.258, longitude: −71.285, altitude: 1175 m a.s.l.). We harvested seeds from 25 individual trees located at a minimum distance of 30 m in order to preclude family relationships. Equal amount of seeds from each mother plant were pooled for the experiments. Seeds were germinated as described in [23] and seedlings were grown in 90 cm^3^ pots in the greenhouse for 2 years prior to the experiments.

Works in angiosperm species such as *A. thaliana*, rice and poplar demonstrate that diurnal cycles of light or temperature affect the expression of a wide proportion of the transcriptome [24–26]. Moreover, in *A. thaliana*, over 75% of heat-responsive transcripts show a time of day-dependent response, and it was demonstrated that both diurnal and circadian regulation of the transcriptome impact experimental interpretation of the heat stress response [27]. With the aim to detect genes regulated by heat stress in *N. pumilio*, and reduce the aforementioned possible diurnal (photocycles-driven) and time of the day-dependent bias in the interpretation of the heat stress experiments, we used the following protocol. Plants were grown for 10 days in growth chambers (SCE BD/600, Bariloche, Argentina) at 20°C with 12 hours light (200 µmol m^−2^ s^−1^, Osram DULUX L 36W) and 12 hours darkness. Then, they plants were subjected to continuous light (100 µmol m^−2^ s^−1^) in order to discard diurnal effects on the regulation of the transcriptome and exposed to two temperature treatments: one group of plants (20 plants per biological replicate) was exposed to 34°C (heat stress), while another 20 plants were kept at 20°C (control). Samples were collected at 48 and 60 hours after the beginning of these temperature treatments. As further explained in the differential expression analysis section, sampling at 48 and 60 hours after the beginning of the temperature treatments allowed us to study common genes up or down-regulated by heat stress at two time points, diminishing the bias of the time of the day on the interpretation of our experiments. Additionally, sampling under continuous light allowed us to discard photocycle-driven effect on the regulation of the transcriptome, allowing us to focus on those genes that were mostly regulated by heat. Each sample consisted of a pool of one whole leaf from 10 seedlings. Samples were immediately frozen in liquid nitrogen and stored at −80°C until the RNA extraction. The experiments were performed twice in the same growth chambers, using different seedlings (two independent biological replicates), yielding a total of 8 samples.

### RNA extraction, library construction and sequencing

Each pool of 10 leaves was manually grounded with mortar and pestle under liquid nitrogen. Total RNA was extracted according to [28], treated with RQ1 RNAse-free DNAse (Promega), and purified with RNeasy Plant Mini Kit (Qiagen), following the manufacturer’s instructions. The integrity of the RNA was assessed in a 0.8% agarose gel, and its quantity and quality with a NanoDrop (ThermoFisher Scientific) and a BioAnalyzer 2100 Plant RNA Pico chip (Agilent) before proceeding with library preparation.

Mature mRNA was selected with Dynabeads mRNA DIRECT Micro Kit (ThermoFisher Scientific), adding ERCC RNA Spike-In Control Mix from the same manufacturer. Eight whole transcriptome libraries were constructed with Ion Total RNA-seq Kit v2 (ionTorrent, Life Technologies), followed by emulsion PCR in an Ion OneTouch 2 System, using the Ion PI Hi-Q OT2 200 Kit (ionTorrent, Life Technologies).

Sequencing was performed using the ionTorrent Proton System (Life Technologies), in a total of three runs (with three, three, and two libraries, respectively) in order to ensure approximately 25 million reads per library, which was shown to be sufficient to detect more than 90% of genes in eukaryotes [29].

### Datasets processing and assembly

Reads were quality-checked with FastQC [30] and trimmed with Trimmomatic [31] (version 0.33; parameters: LEADING:3 TRAILING:3 SLIDINGWINDOW:5:15 MINLEN:36) and the Fastx toolkit [32] trimmer (version 0.0.13; parameters: −Q33 −l 250).

Trimmed reads were assembled using SPAdes [33] (version 3.11.0; parameters: ╌rna ╌iontorrent −k67 ╌ss-fr). The final *k* −mer value of 67 was chosen after several assemblies with different *k* −mer values (five in total, between 21 and 77). The assembled contigs that overlapped considerably were expanded, and highly redundant contigs were eliminated.

After assembly, trimmed reads were mapped back to the assembly using STAR [34] (version 2.4.2a; genome indexing parameters: ╌runMode genomeGenerate ╌genomeSAindexNbases 11; mapping parameters: ╌outSAMunmapped Within ╌alignIntronMax 21 ╌outFilterScoreMinOverLread 0.4 –outFilterMatchNminOverLread 0.4) as a measure for the percentage of reads used in the assembly. In order to assess the functional completeness of the new reference assembly, 248 Core Eukaryotic Genes models [35] and 2121 eudicotyledon single-copy orthologs (BUSCO; [36]) were run against the assembly. Finally, *N. pumilio* Sanger sequences available from GenBank were searched in the assembly to check for completeness and sequence identity.

### Annotation

The transcriptome assembly was annotated against the *Arabidopsis thaliana* proteome (http://www.uniprot.org/proteomes/UP000006548), using the longest ORF per frame per contig as query. We chose this species because of its long-standing status as a plant model species, being used as a reference for annotation of other species, such as the model tree *Populus trichocarpa*, and because of the many resources available online, such as expression atlases under diverse stress conditions (AtGenExpress from http://www.arabidopsis.org), or circadian time series expression curves (http://diurnal.mocklerlab.org). Contigs that were not annotated against the *A. thaliana* proteome were in turn annotated against SwissProt (https://www.uniprot.org/uniprot/?query=reviewed:yes). For annotation, contigs were aligned to the database (*A. thaliana* proteome) using BLAT [37], and a file with a single best-hit annotation for each successful contig was generated. The annotation file features Gene Ontology (GO; [38]) terms for each annotated contig from the corresponding *A. thaliana* subject (GO terms downloaded from Gene Ontology Annotation Database, https://www.ebi.ac.uk/GOA).

### Differential expression analysis

Reads from all libraries were quantified against the reference assembly using Salmon, version 0.8.1 [39]. After quantification, a tab-delimited file containing the unnormalized expression level for each contig in each of the eight libraries was put together. For differential expression analysis, DESeq2 [40] was used in an R environment, with default models and parameters. The two temperatures (20°C and 34°C) were contrasted, taking the different moments of the day as biological replicates; that is, four biological replicates for each temperature were compared. This protocol allowed us to reduce the bias of the time of the day on the interpretation of our experiments, focusing our study in the detection of genes that were particularly induced by heat stress. Contigs with an FDR*<*0.05 were considered as differentially expressed between temperatures.

### Functional enrichment analysis

Combining the output table from DESeq2 with the annotations produced for the assembly, we were able to perform GO terms and metabolic pathways enrichment in contigs over- and under-expressed in response to temperature. For GO term enrichment analysis, PANTHER version 14.1 [41] was used via its implementation in the TAIR (The Arabidopsis Information Resource; https://www.arabidopsis.org/) website. For KEGG (Kyoto Encyclopedia of Genes and Genomes; [42]) metabolic pathways enrichment analysis, KOBAS 3.0 online tool was used [43]. In both cases, the background dataset were all *A. thaliana* identifiers present in our assembly’s annotation. Visualization and clustering of over-represented GO terms was performed with REVIGO [44].

The annotated transcription factors were classified into their corresponding families using the Plant Transcription Factor Database [45] gene annotation for *A. thaliana* (http://planttfdb.gao-lab.org/index.php?sp=Ath). A Fisher exact test was carried out for each family and each temperature treatment, and families were sorted from most to less enriched at each temperature. For functional regulatory analysis, PlantRegMap [46] regulation prediction tool (http://plantregmap.gao-lab.org/regulation_prediction.php) was run on over-expressed genes.

In order to inspect shared molecular components that are up- or down-regulated in response to heat stress among plant species, we used expression data from *A. thaliana* [47] *and Populus tomentosa* [18]. These papers were selected among those published in recent years because they feature a complete, publicly available set of gene annotations and differential expression statistical results. The *P. tomentosa* study consisted in RNA-seq experiments where contigs were annotated against *P. trichocarpa*, a related species with a sequenced genome, which in turn was annotated against *A. thaliana*. Thus, we were able to obtain corresponding *A. thaliana* IDs for annotated contigs for all three species that could be intersected and provided us with a list of shared genes in the three species. Among these, each species had a set of differentially expressed genes, which were also intersected and subjected to GO term enrichment and transcription factor regulation prediction as described above for *N. pumilio*.

### Primer design and quantitative RT-PCR validation

In order to evaluate the accuracy of our transcriptome data, a total of 13 (eight up-regulated and five repressed in response to high temperature) genes were selected to carry out a qRT-PCR analysis. We chose a group of contigs that allowed to test a wide range of expression (from intermediate to high expression; between 5 and 45 Transcripts Per Million averaged across conditions) and fold change (Log2 fold change between −11 and 8) in our RNA-seq data. Primers were designed with Primer-BLAST [48] (S1 Table).

RNA was extracted and purified using the aforementioned protocols, from leaf samples of two independent experiments performed in the same chambers and conditions as those that were used to produce the transcriptomic libraries. cDNA synthesis was performed using M-MLV Reverse Transcriptase and RNasin Ribonuclease Inhibitor (Promega), and quantitative PCR reactions were done using SsoAdvanced Universal SYBR Green Supermix (Bio-Rad) in a CFX96 Touch device (Bio-Rad) according to manufacturer’s instructions. Each qPCR reaction consisted of three technical replicates. For relative gene expression analysis, two reference genes (*DER2*.*2* and *P2C22*; [49]) and the ΔΔ*C*_*t*_ [50] method were used.

### Code availability

The scripts used for assembly improving and annotation are openly available through GitHub [51] via an MIT License.

## Results

### RNA sequencing, *de novo* transcriptome assembly and gene annotation

A total of 222,828,783 reads were sequenced for the eight libraries (Table 1). The read throughput and average length were in accordance with the device specifications [52]. Raw reads were trimmed to eliminate low-quality ends, and low-quality sequences (Q*<*20) were removed. This procedure allowed us to increase the overall read quality at the expense of shorter overall read length (Table 1). Pearson’s correlation tests showed high levels of reproducibility among the biological replicates, with R values ranging between 0.86 and 0.90 (S1 Fig).

**Table 1.**
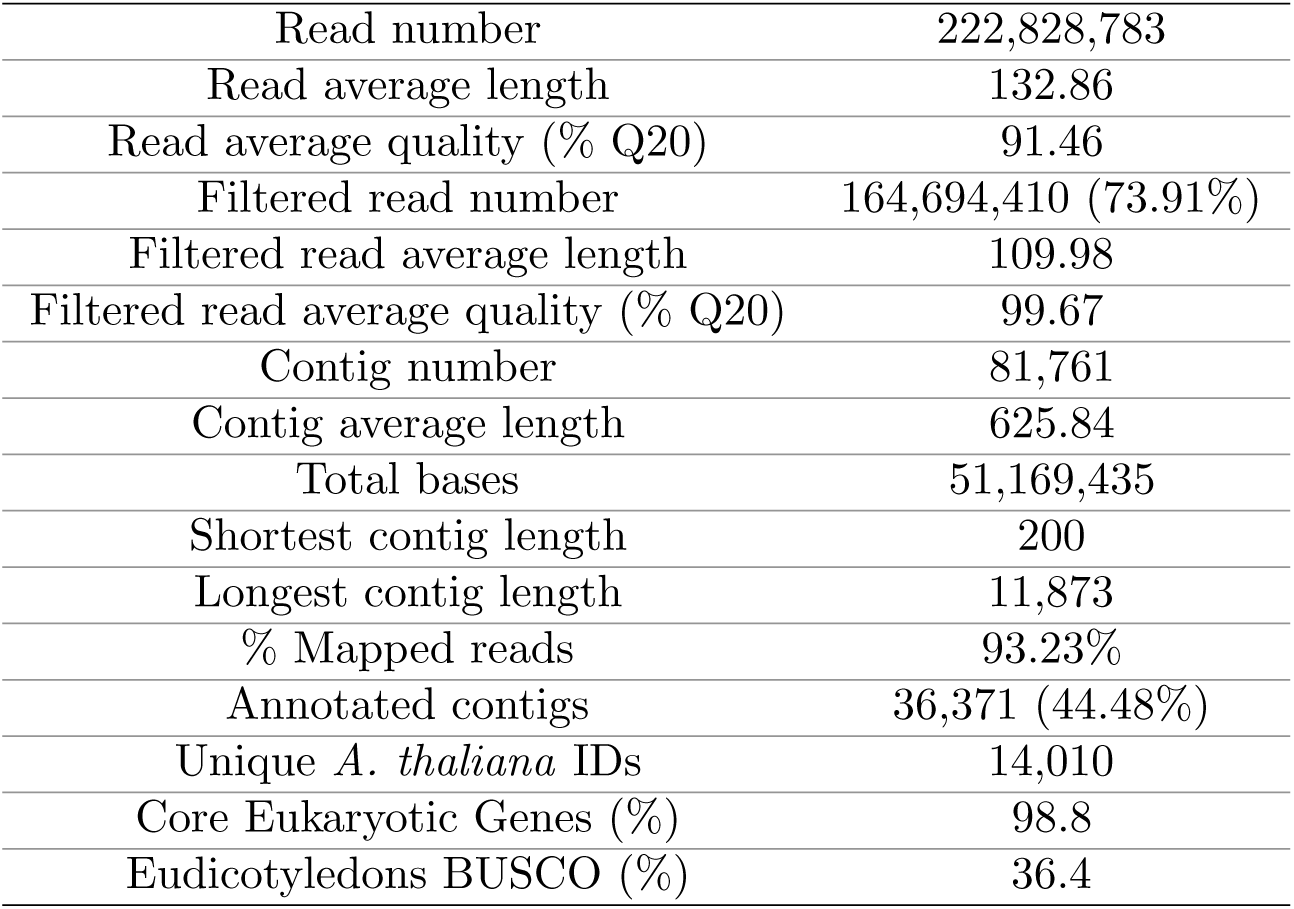
*Nothofagus pumilio* transcriptome statistics.

|  |  |
| --- | --- |
| Read number | 222,828,783 |
| Read average length | 132.86 |
| Read average quality (% Q20) | 91.46 |
| Filtered read number | 164,694,410 (73.91%) |
| Filtered read average length | 109.98 |
| Filtered read average quality (% Q20) | 99.67 |
| Contig number | 81,761 |
| Contig average length | 625.84 |
| Total bases | 51,169,435 |
| Shortest contig length | 200 |
| Longest contig length | 11,873 |
| % Mapped reads | 93.23% |
| Annotated contigs | 36,371 (44.48%) |
| Unique *A. thaliana* IDs | 14,010 |
| Core Eukaryotic Genes (%) | 98.8 |
| Eudicotyledons BUSCO (%) | 36.4 |

In order to capture a complete and non-redundant set of genes to perform a differential expression analysis, a single transcriptome was assembled using the eight sequenced libraries. Contigs that overlapped considerably but were assembled as separate contigs by SPAdes were merged [51], thus yielding a total of 81,761 contigs, with a read utilization of 93.23% (Table 1). Almost half of all contigs (44.48%) were annotated against the *A. thaliana* UniProt proteome, whereas just a small proportion of the remaining contigs (6.4% of those non-annotated, which means 3.5% of all contigs) were annotated against SwissProt. Thus, we worked with *A. thaliana* UniProt database for transcriptome annotations. Most of the annotated contigs (83%) were longer than 500 nts, whereas 92% of the non-annotated contigs were shorter than 500 nts (S2 Fig). Regarding functional completeness, 245 (98.8%) Core Eukaryotic Genes models and 772 (36.4%) eudicotyledon BUSCOs were found in the *N. pumilio* assembly. Alignment of available (N=5) published Sanger sequences from *N. pumilio* to our transcriptome yielded more than 98% match, further supporting the high quality of the *de novo* assembled contigs (S2 Table).

Each sample was submitted as a BioSample in the NCBI BioProject PRJNA414196, and the raw sequences were deposited in the NCBI Sequence Read Archive (SRA). The assembly was deposited in the NCBI Transcriptome Shotgun Assembly (TSA) database (S3 Table). This reference transcriptome was then used for differential expression and downstream analyses.

### Differential expression analysis in response to heat

The four libraries from the heat and control treatment respectively were taken as replicates in the differential expression analysis. A total of 5,214 contigs were found to be differentially expressed between temperatures (FDR*<*0.05; 6.38% of all assembled contigs). Of these, 3,358 (64.4% of differentially expressed contigs) were up-regulated and 1,856 (35.6%) were repressed in response to the heat treatment (S3 Fig). Out of these 1,633 of the up-regulated and 1,345 of the repressed contigs could be annotated and accounted for 1,265 and 883 unique protein IDs, respectively (S13 Table and S14 Table).

### Pathways and biological processes promoted and repressed by heat

The protein IDs from *A. thaliana* corresponding to the annotated, differentially expressed contigs were evaluated for KEGG pathway and GO term enrichment for each group separately (up-regulated and repressed in response to high temperature).

A total of 23 and 40 KEGG pathways were enriched in genes repressed or promoted at 34°C, respectively. Most of the pathways exclusively over-represented in genes repressed at 34°C (Fig 1A, S4 Table) were directly related to photosynthesis, for example “Photosynthesis”, “Photosynthesis – antenna proteins”, “Carotenoid biosynthesis”, or “Porphyrin and chlorophyll metabolism”. Other pathways related to basic cell metabolism were enriched in both promoted and repressed groups of genes, but more so in the group repressed at 34°C (Fig 1C). These pathways included “Carbon fixation in photosynthetic organisms”, “Glyoxylate and dicarboxylate metabolism”, “Pentose phosphate pathway”, “Carbon metabolism” and “Nitrogen metabolism”. In contrast, enriched pathways in genes over-expressed at 34°C were mostly related to stress responses like “Biosynthesis of secondary metabolites” (Fig 1C). Among the various families of secondary metabolites, many enriched pathways exclusively present in genes promoted by 34°C (Fig 1B) were specific to the biosynthesis of stress-related metabolites families such as flavonoids, phenylpropanoids, mono-, sesqui-and tri-terpenoids, and some groups of alkaloids. Pathways related to translation and protein processing were also triggered at 34°C, as indicated by several enriched pathways such as “Ribosome”, “Protein processing in endoplasmic reticulum”, “Spliceosome” or “RNA transport” (Fig 1B and C, S5 Table).

**Fig 1.**
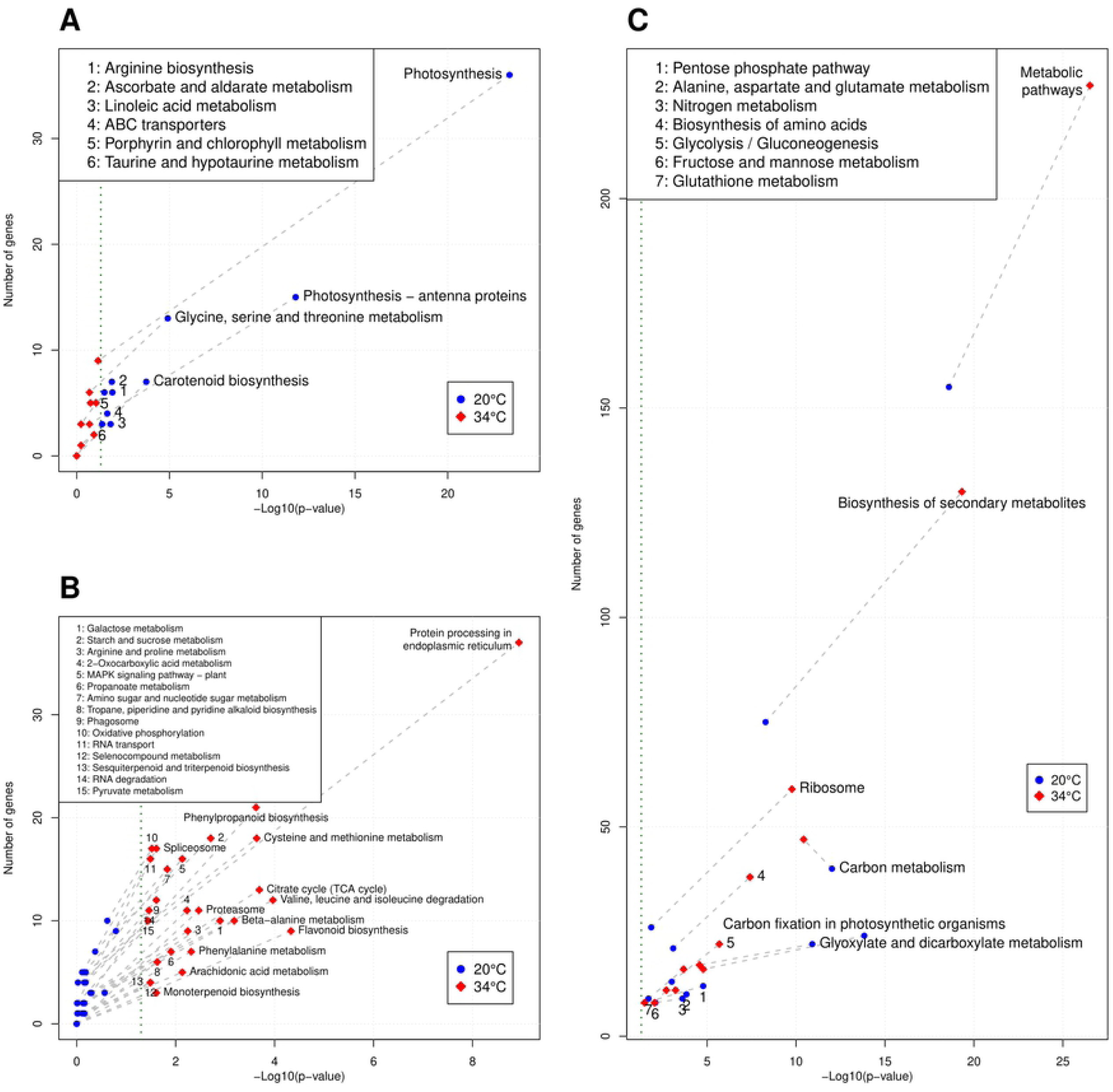
KEGG enriched pathways. **A**: Pathways significantly enriched at 20°C but not enriched at 34°C. **B**: Pathways significantly enriched at 34°C but not enriched at 20°C. Vertical green line indicates *p* = 0.05. **C**: Pathways enriched at both temperatures.

Overall, a total of 78 GO terms were enriched in genes repressed at 34°C (S6 Table) and 71 GO terms were enriched in genes induced by 34°C (S7 Table). Semantic reduction and clustering of enriched GO biological processes show that “photosynthesis”, “glucose metabolism” and “generation of precursor metabolites and energy” were the main processes repressed by heat, whereas at 34°C the response to various stress signals such as “response to chemical” or “secondary metabolism” and “protein folding / refolding” were highly significant (Fig 2). Moreover, the most enriched GO terms in genes repressed at 34°C in all three branches of the ontology (Biological Process, Molecular Function and Cellular Component) were related to photosynthesis, whereas among the genes up-regulated at 34°C these terms corresponded to specific stresses together with those related with translation, ribosome activity and protein processing (Fig 2, S4 Fig and S5 Fig). This indicates the high coherence and complementarity between GO and KEGG enrichment analyses. Furthermore, the response to misfolded or topologically incorrect proteins and their degradation via proteasome were up-regulated at 34°C (S5 Table and S7 Table). Tables 2 and 3 show all chaperones and ubiquitin-ligases significantly more expressed at 34°C. The great number and diversity of these proteins suggests the importance of protein re-folding and degradation in the response to high temperature stress.

**Fig 2.**
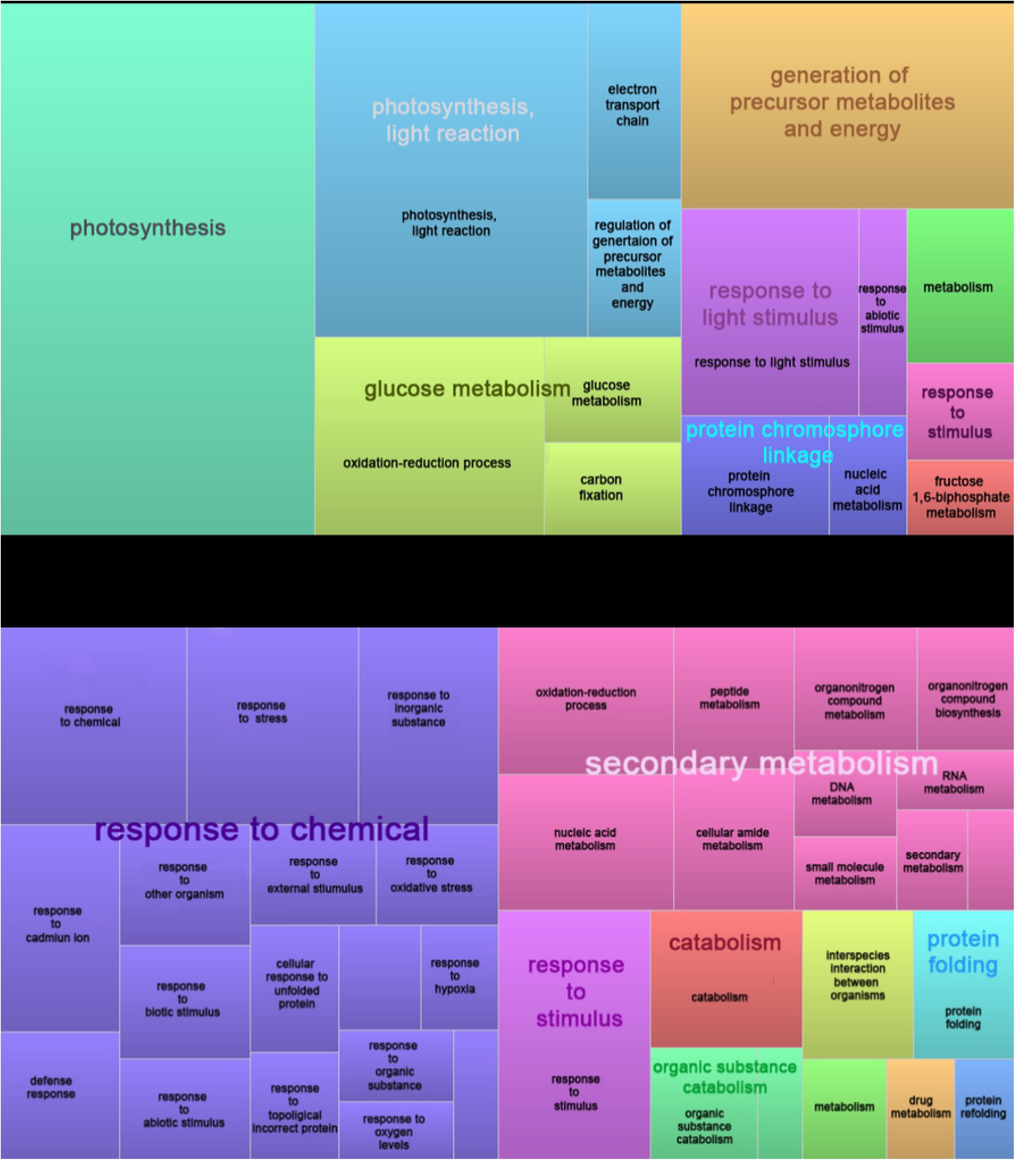
**Semantically reduced overrepresented Gene Ontology biological processes in genes repressed (A) and promoted (B) in response to high temperature**

**Table 2.** **Heat Shock Proteins (HSPs), Late embryogenesis abundant (LEAs) and Dehydrins significantly more expressed at 34°C**

| Family | Name | *A. thaliana* UniProt ID |
| --- | --- | --- |
| HSP | 15.7 kDa heat shock protein | Q9FHQ3 |
|  | 17.6 kDa class I heat shock protein 1 | Q9XIE3 |
|  | 17.6 kDa class I heat shock protein 2 | Q9ZW31 |
|  | 18.1 kDa class I heat shock protein | P19037 |
|  | 22.0 kDa heat shock protein | Q38806 |
|  | 23.6 kDa heat shock protein | Q96331 |
|  | Heat shock 70 kDa protein 5 | Q9S9N1 |
|  | Heat shock 70 kDa protein 6 | Q9STW6 |
|  | Heat shock 70 kDa protein 8 | Q9SKY8 |
|  | Heat shock 70 kDa protein 9 | Q8GUM2 |
|  | Heat shock 70 kDa protein 10 | Q9LDZ0 |
|  | Heat shock 70 kDa protein 18 | Q9C7X7 |
|  | Heat shock protein 21 | P31170 |
|  | Heat shock protein 90-1 | P27323 |
|  | Heat shock protein 90-2 | P55737 |
|  | Heat shock protein 90-6 | F4JFN3 |
| LEA | Late embryogenesis abundant protein 2 | Q9SRX6 |
|  | Late embryogenesis abundant protein 3 | Q9SA57 |
|  | Late embryogenesis abundant protein 41 | Q39084 |
|  | Late embryogenesis abundant protein 46 | Q9FG31 |
|  | Late embryogenesis abundant protein family protein | F4IYB7 |
| Dehydrins | Dehydrin Xero 1 | P25863 |
|  | Dehydrin Rab18 | P30185 |

**Table 3.** Ubiquitin-ligases significantly more expressed at 34°C

| Name | *A. thaliana* UniProt ID |
| --- | --- |
| E3 ubiquitin-protein ligase | Q9LHE6 |
| E3 ubiquitin-protein ligase ATL41 | Q9SLC3 |
| E3 ubiquitin-protein ligase UPL2 | Q8H0T4 |
| E3 ubiquitin-protein ligase UPL3 | Q6WWW4 |
| E3 ubiquitin-protein ligase UPL4 | Q9LYZ7 |
| E3 ubiquitin-protein ligase UPL5 | Q9SU29 |
| E3 ubiquitin-protein ligase PUB22 | Q9SVC6 |
| E3 ubiquitin-protein ligase PUB23 | Q84TG3 |
| E3 ubiquitin-protein ligase RZFP34 | Q9FFB6 |
| E3 ubiquitin protein ligase DRIP2 | Q94AY3 |
| Probable E3 ubiquitin-protein ligase EDA40 | F4JSV3 |

### Regulation of transcription and hormone signaling in heat stress

A total of 703 TFs belonging to 55 families were annotated in the transcriptome of *N. pumilio*. Of these, 59 (from 20 families) were over-expressed at 34°C and 20 (from 15 families) were repressed at 34°C (S8 Table). Families that showed a higher representation among genes repressed by heat stress include MYB-like, ARF (Auxin Response Factor), CAMTA (Calmodulin-binding Transcription Factor) and RAV (Related to ABI3/VP1). On the contrary, members of families like MYB, ERF (Ethylene Response Factors), HSF (Heat Stress Factor), NAC (NAM, ATAF1/2 and CUC2), WRKY, WOX (WUSCHEL-related homeobox), LBD (Lateral Organ Boundaries Domain), and EIL (Ethylene-Insensitive 3-like) were among the TF families that showed a bias towards up-regulation in response to heat. A total of 137 TFs were found to have over-represented targets among the genes over-expressed at 34°C (S9 Table). Among these TFs there were representatives of families up-regulated by heat such as MYB, ERF, NAC, and WRKY families, apart from others like ZAT proteins or NLP4.

Hormones play a fundamental role in plant stress responses, and the function of particular hormones and their crosstalk differ among tree species [3]. The homolog of AHP5 (Histidine-containing phosphotransfer protein 5), an important two-component mediator between cytokinin sensing and its response regulators, was repressed at 34°C (S13 Table). In addition, ARFs TF family was down-regulated at 34°C (S8 Table). On the contrary, 34°C promotes the accumulation of the *N. pumilio* homolog of EIN3 (Ethylene-insensitive 3) and several ERF TFs, indicating that ethylene signaling and response are promoted by heat stress. In addition, 4 out of 7 ABA phosphatases belonging to the clade A [53] are over-expressed in *N. pumilio* in response to heat stress (S14 Table). These ABA phosphatases, which show high homology to the ABA phosphatases At4g26080, At5g59220, At1g07430 and At3g11410 of *A. thaliana*, are part of the KEGG pathway “MAPK signaling pathway – plant”, over-represented in genes induced by 34°C (Fig 1B).

### Comparative heat stress responses between species

The comparative analysis of heat stress response of *N. pumilio, A. thaliana* and *P. tomentosa*) resulted in 68 genes that were significantly more expressed at high temperature in all three species (S10 Table), many of them belonging to the aforementioned up-regulated groups in *N. pumilio*, such as HSPs, LEAs (Table 2), and HSF and WRKY TF families. In addition, one common gene was a constituent of the large ribosomal protein (60S), and 7 out of the 68 genes were involved in the “Spliceosome” pathway. These results suggest the existence of conserved cores of regulation of gene expression in response to stress at transcriptional and translational levels in angiosperms. In addition, common genes included proteins involved in the regulation of protein folding in the endoplasmic reticulum lumen, such as Derlin-1 and the DnaJ protein ERDJ3A or the DNAJ protein P58IPK homolog, that contribute to the protection of cells to endoplasmic reticulum stress [54].

GO enrichment analysis showed that the main processes shared among species in response to heat were those related to protein misfolding and refolding, apart from general and specific stress terms (S11 Table and S6 Fig). Moreover, a total of 52 common TFs were found to have enriched targets among the genes over-expressed at high temperature (S12 Table). Of those, 29 (55%) were ERFs, demonstrating the relevant role of ethylene in the response to high temperature stress in these species.

### Validation of RNA-seq data with quantitative RT-PCR

To verify the validity of our RNA-seq differential expression results, we analysed the expression of 13 genes by quantitative RT-PCR (eight up-regulated and five repressed in response to high temperature). The correlation between the gene expression values for the two methods was high (R^2^=0.793; Fig 3) and the expression trends were consistent. These results show the high reliability of the RNA-seq data.

**Fig 3.**
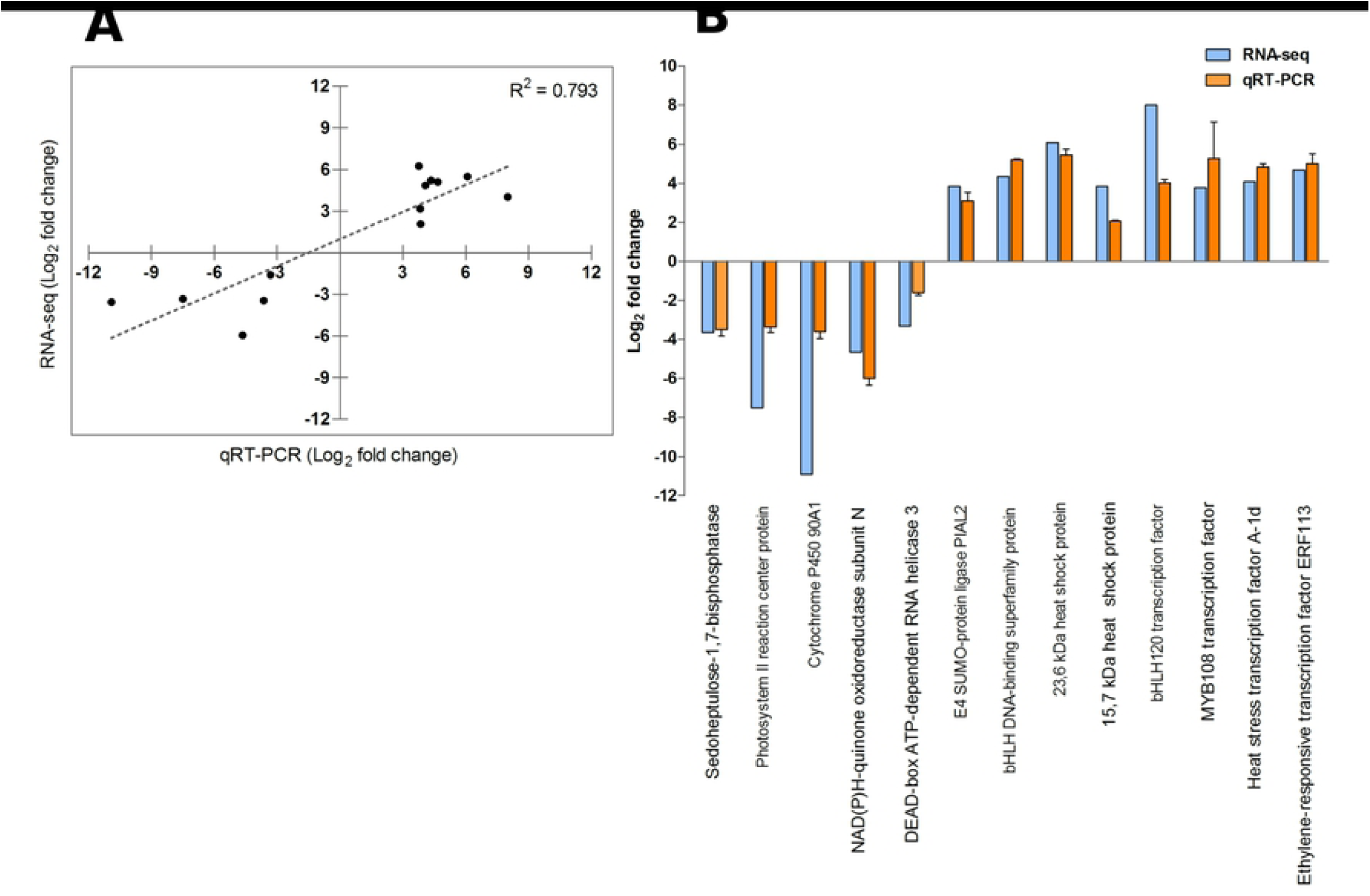
Verification of 13 differentially expressed genes by qRT-PCR. **A**: Pearson linear correlation. **B**: Bar plot. Error bars represent SD of 3 technical replicates

## Discussion

Through the assembly of the first transcriptome and the performance of RNA-seq analyses of *Nothofagus pumilio*, we identified the main molecular and biological pathways affected during heat stress in this tree species. In addition, by analyzing overlapping up-regulated genes in experiments of heat stress in *N. pumilio, P. tomentosa* and *A. thaliana* we identified common candidate genes for heat stress response across angiosperm species with potential biotechnological applications.

After assembling, annotating and analyzing the transcriptome for *N. pumilio*, we identified 5,214 differentially expressed contigs in response to heat. Interestingly, the number of up-regulated contigs was almost twice the down-regulated ones (S3 Fig). This suggests that the heat response in *N. pumilio* involves the rearrangement of a relevant fraction of its transcriptome, and is characterized by the induction, rather than the repression, of the expression of a large proportion of genes. These findings contrast with transcriptomic studies under heat stress in other tree species such as *Populus tomentosa, P. simonii* and *Abies koreana*, where a balanced proportion of contigs was down vs. up-regulated [18], or even the proportion of down-regulated contigs at warm temperatures was larger than the up-regulated ones [15, 17]. The high correlation in gene expression values between *in silico* analysis and qRT-PCR experiments (Fig 3) showed the reliability of our RNA-seq data.

Functional enrichment allowed us to identify the main underlying pathways and biological processes affected in the transcriptome of *N. pumilio* in response to heat stress. In accordance with reports from other tree species such as *Olea europaea, Quercus lobata, Pseudotsuga menziesii, Pyrus betulaefolia, Camellia sinensis*, and *Santalum album* under cold or drought stress [13, 55–59], genes repressed in response to heat in *N. pumilio* showed an over-representation of KEGG pathways and GO terms related to photosynthesis as a whole. These genes were also implicated in sub-processes like biosynthesis of primary (chlorophyll) and auxiliary (carotenoid) photosynthetic pigments, or the action of photosynthesis antenna proteins. Similarly, carbon metabolism was over-represented as a whole in the repressed genes, and so were several processes like the metabolism and biosynthesis of simple sugars (glucose, hexoses, monosaccharides), or the biosynthesis of fatty acids through the glyoxylate cycle and linoleic acid metabolism. In contrast, the analysis of the up-regulated genes indicated that heat triggers an abrupt adjustment of translation, as evidenced by the over-representation of KEGG pathways and GO terms related to protein processing, the ribosome, peptide biosynthesis, and translation (Fig 1, Fig 2). A well-known effect of abiotic stress is the production of Reactive Oxygen Species (ROS), which can oxidize biomolecules and set off cell death [60]. Many plant secondary metabolites have antioxidant properties, and their production is significantly increased by abiotic stress [60, 61]. In our study, genes involved in the biosynthesis of many antioxidant metabolites families were found to be triggered by high temperature, namely phenylpropanoids, flavonoids, mono-, sesqui- and tri-terpenoids, and tropane, piperidine and pyridine alkaloids. Moreover, it has been shown in trees that MAPK cascades promote antioxidant responses [62], and the plant MAPK signaling pathway was enriched in genes more expressed at 34°C in *N. pumilio* (Fig 1B and S5 Table).

Our analysis indicated that the response to misfolded or topologically incorrect proteins was up-regulated by heat stress. In plants, HSPs and other chaperones bind to misfolded proteins, which are in turn ubiquitinated and directed to the proteasome for their degradation [63, 64]. Several chaperones (including HSPs) and ubiquitin-ligases were found to be significantly more expressed at 34°C than at 20°C (Tables 2 and 3), indicating the importance of these processes in the response to high temperature stress. In concordance, forestry species such as *Quercus lobata, Pseudotsuga menziesii* and *Prunus persica* show over-expression of chaperones and ubiquitin-ligase proteins in response to different abiotic stresses [13, 56, 65], indicating that the induction of protein re-folding and ubiquitination followed by degradation by the proteasome pathway constitutes a relevant molecular strategy that allows trees to cope with adverse abiotic conditions.

Transcription Factors (TFs) are known to play important roles in the transcriptional regulation of stress responses, and their involvement in many biotic and abiotic stresses across plant species has been extensively reviewed [53, 66, 67]. In *N. pumilio*, TFs belonging to families such as WRKY, WOX, LBD and NAC were positively regulated in response to high temperature (S8 Table). In concordance, previous reports show that NAC TFs play roles in numerous biotic and abiotic stresses including heat [68, 69], and whereas WOX TFs promotes the response to abiotic stresses such as drought and salinity in *Brassica napus* [70, 71], LBD TFs were suggested to play roles in the response to cold in *Broussonetia papyrifera* [72]. The WRKY gene family is one of the largest TF family in plants, playing roles in the regulation of a broad range of physiological and developmental processes [73], including the response to biotic and abiotic stress [74, 75]. It is interesting to note that most of the WRKY TFs identified in *N. pumilio* over-expressed at 34°C have homologs that are involved in the response to abiotic stress in *Arabidopsis*, rice or poplar. For example, WRKY17, 45 and 53 were shown to participate in the response to drought of rice and *Arabidopsis* [76–79], and WRKY75 is involved in the response to salt stress in poplar trees [80]. Moreover, WRKY18, 48 and 53 are induced by ROS in *Arabidopsis* [81, 82]. This is in accordance with the over-representation of KEGG pathways and GO terms related to response to oxidative stress at 34°C (Fig 1, Fig 2), raising the hypothesis that ROS may induce the expression of a sub-set of WRKY TFs during the response to high temperature in *N. pumilio*. Genes up-regulated at 34°C show an enrichment of targets of WRKY TFs (S9 Table), further supporting the proposed relevance of WRKY TFs in the modulation of the response to heat of *N. pumilio*. In contrast, the homolog of RAV1, which was shown to negatively regulate drought and salt stress responses independently of ABA in *Arabidopsis* [83], was strongly repressed by heat stress in the *N. pumilio* transcriptome (S13 Table).

In relation to hormonal signaling, ethylene is an important plant hormone which is known to be involved in stress responses [84]. Several members of the ERF family were over-expressed under heat stress in *N. pumilio* (S8 Table and S14 Table), and genes up-regulated at 34°C showed an enrichment of ERF targets (S9 Table). Moreover, the homolog of EIN3, a master regulator of ethylene signaling [85], together with several of its targets [86], were over-expressed in heat-treated plants (S14 Table), indicating that the ethylene pathway is activated at 34°C. Additionally, our data suggest that ABA signaling and response constituted another hormonal pathway up-regulated by heat. This is supported by the over-expression of several ABA-responsive genes such as those described in [53], including dehydrins, LEAs and protein phosphatases of the clade A in plants exposed to high temperature (Table 2 and S14 Table). Furthermore, genes over-expressed at 34°C showed enriched targets of TFs related to the regulation of ABA signaling (S9 Table). In contrast, our data indicates that auxin signaling and re-localization is repressed in response to heat in *N. pumilio*. This is evidenced by the fact that ARFs, which are relevant components of the auxin signaling pathway [87], were repressed by high temperature (S8 Table). In addition, auxin-efflux ABC transporters were down-regulated under heat stress (Fig 1A). Finally, RVE1 (Reveille 1), a MYB-like TF which links the circadian clock with the auxin signaling pathway [88], was down-regulated by high temperature (S13 Table).

Most of our knowledge on the molecular bases of heat stress was originated from studies focused on a single species, and comparisons between two or more species regarding their common or distinct response mechanisms are scarce. In this study, the combined analysis of genes over-expressed under heat stress in *N. pumilio, A. thaliana* and *P. tomentosa* allowed us to identify a core of shared responses to high temperature, mostly related to protein misfolding and chaperone activity, with the over-expression of more than ten HSPs, one LEA and two DnaJ protein genes (S10 Table, S11 Table, and S6 Fig). Alternative splicing (AS) is known to be triggered in plants in response to stress [89], and particularly, several genes related to plant stress responses are subjected to AS [90]. In our study, the “Spliceosome” KEGG Pathway was significantly enriched among genes up-regulated at 34°C in *N. pumilio* (Fig 1B and S5 Table), and several genes involved in pre-mRNA splicing were shared between *N. pumilio, A. thaliana* and *P. tomentosa* at high temperature, including several DEAD-box ATP-dependent RNA helicases (S10 Table). These results support the fact that AS constitutes an important mechanism in plant response to abiotic stress and highlight the potential relevance of a subset of genes associated with the splicing machinery in the response to heat stress across angiosperms.

Regarding hormone signaling, many ERFs were found to have enriched targets among genes over-expressed at high temperature in the three species (S12 Table), indicating the relevance of ethylene and ERF TFs in the response to heat, and further supporting the reported results in *N. pumilio*. Apart from ERFs, our analysis allowed us to identify common targets or relevant TF families already discussed such as WRKY and NAC (S12 Table), and one of the TFs with most significantly enriched targets, and the single most enriched considering only *N. pumilio* (S9 Table) was NLP4, a member of the NLP (NIN-like Protein) family. Members of this family have been recently shown to be differentially expressed in response to cold, heat and drought treatments in rice [91]. Finally, the zinc finger protein ZAT10, which constitutes a transcriptional repressor involved in abiotic stress responses [92], was over-expressed under heat stress in the three species (S10 Table), and *A. thaliana, P. tomentosa* and *N. pumilio* transcriptomes of heat-treated plants show an enrichment of ZAT10 targets (S12 Table). This suggests that the ZAT10 regulon constitutes a relevant regulatory module during heat stress responses in angiosperms. All these results suggest a strong shared core of transcriptional and translational regulation of gene expression in response to abiotic stress in plant species of potential biotechnological application.

## Conclusion

This work constitutes the first report on whole transcriptome analysis in the *Nothofagus* genus. Through RNA-sequencing and bioinformatic analysis, we were able to identify a wide spectrum of heat-responsive transcripts, including 59 transcription factors, and revealed several features of the molecular adjustment strategy of *N. pumilio* to heat stress. The down-regulation of photosynthesis and sugar metabolism, together with the promotion of the expression of stress response genes are indicative of a trade-off between growth and survival, and suggest that carbon sequestration can be severely affected in *N. pumilio* in a context of global warming. Our data provide evidences of the prominent role of WRKY TFs in the response to heat in *N. pumilio*, not previously highlighted in other studies in tress. The evidenced up-regulation of ethylene and ABA pathways and the repression of auxin signaling and re-localization in response to high temperature are indicative of a complex transcriptional landscape with highly variable interactions and cross-talk between hormone signal transduction pathways.

Furthermore, the enrichment of biological pathways related to the spliceosome, protein ubiquitination and MAP kinase cascades suggests that heat stress in *N. pumilio* is governed by a multi-layered, fine-tuned regulation of gene expression. The identification of overlapping genes up-regulated under high temperature in *N. pumilio, P. tomentosa* and *A. thaliana* provides candidates for engineering plants in order to promote heat stress resistance. Thus, this study represents an important step towards the possibility of breeding acceleration, genomic markers development, genotype selection and *in vivo* risk assessment for *N. pumilio* with potential use in other plant species.

## Supporting information

**S1 Table Primer sequences for qRT-PCR validation**.

**S2 Table Comparison between assembled NGS and Sanger *Nothofagus pumilio* sequences**.

**S3 Table *Nothofagus pumilio* transcriptome deposition information**.

**S4 Table Overrepresented pathways in genes repressed at 34**°**C**

**S5 Table Overrepresented pathways in genes promoted at 34**°**C**

**S6 Table Overrepresented GO terms in genes repressed at 34**°**C**

**S7 Table Overrepresented GO terms in genes promoted at 34**°**C**

**S8 Table Transcription factor families annotated in *N. pumilio* transcriptome**. An asterisk indicates an enrichment ratio larger than 2, i.e. more than twice TFs observed than expected for the corresponding temperature.

**S9 Table Transcription factors whose targets are enriched in genes promoted at 34**°**C**

**S10 Table Genes promoted by high temperature in *N. pumilio, A. thaliana* and *P. tomentosa***.

**S11 Table Overrepresented GO terms in genes promoted by high temperature in *N. pumilio, A. thaliana* and *P. tomentosa***.

**S12 Table Transcription factors whose targets are enriched in genes promoted by high temperature in *N. pumilio, A. thaliana* and *P. tomentosa***.

**S13 Table Annotated contigs repressed at 34**°**C in *Nothofagus pumilio*. S14 Table Annotated contigs promoted at 34**°**C in *Nothofagus pumilio***.

**S1 Fig Pearson’s correlation test between biological replicates. A**: 20°C, 48 hours after onset of temperature treatment (h.a.t.). **B**: 20°C, 60 h.a.t. **C**: 34°C, 48 h.a.t. **D**: 34°C, 60 h.a.t. Low TPM (transcripts per million) values are represented together at the lower end of both axes for better visualization.

**S2 Fig Length distribution of *Nothofagus pumilio* assembled transcripts**. Each bar discriminates between annotated and unannotated contigs of the corresponding length interval.

**S3 Fig Differentially expressed contigs. A**: MA-plot. **B**: Volcano plot. **Red** and **Blue**: differentially expressed contigs. **Black**: contigs not differentially expressed. Total contigs: 81761.

**S4 Fig Semantically reduced overrepresented Gene Ontology molecular functions in genes repressed (A) and promoted (B) in response to high temperature**.

**S5 Fig Semantically reduced overrepresented Gene Ontology cellular components in genes repressed (A) and promoted (B) in response to high temperature**.

**S6 Fig Semantically reduced overrepresented Gene Ontology biological processes (A), molecular functions (B), and cellular components (C) in genes promoted by high temperature in *N. pumilio, A. thaliana* and *P. tomentosa***.

## Acknowledgments

M.E-B. is a postdoctoral fellow from Consejo Nacional de Investigaciones Científicas y Técnicas (CONICET). The authors wish to thank Dr. Erwan Guichoux and all the Genome Transcriptome Facility (INRA, Bordeaux, France) for valuable help in sample processing and library preparation. We also thank the Condensed Matter Theory Group (Centro Atómico Bariloche, Comisión Nacional de Energía Atómica, Bariloche, Argentina) for providing computing facilities.

